# Three photon microscopy of mouse brain structure and function at 2 mm depth and beyond

**DOI:** 10.64898/2026.04.06.716666

**Authors:** Monzilur Rahman, Chi Liu, Dimitre G Ouzounov, Chris Xu

**Affiliations:** School of Applied and Engineering Physics, Cornell University, Ithaca, New York, USA

## Abstract

High-resolution noninvasive imaging of neuronal activity at single-cell resolution deep within brain tissue is essential for understanding brain function and disease. Here we show that an improved 1300-nm three-photon microscope for maximal excitation and collection efficiency enables imaging up to the three-photon depth limit in the intact mouse brain. Our platform achieves structural imaging of brain vasculature at depths up to 2.5 mm and functional imaging of neural activities at depths up to 2 mm, reaching deep brain regions previously inaccessible by multiphoton imaging. These advances extend the frontier of deep-tissue functional imaging and open new possibilities for longitudinal and mechanistic studies in neuroscience and beyond.

## Main

Multiphoton microscopy (MPM) enabled direct observation of cellular activity deep within the living brain ^1–11^. The depth limit of MPM, however, is dictated by the effective attenuation length (EAL), a parameter combining contributions from tissue scattering and absorption, and the signal-to-background ratio (SBR) ^8,12–15^. As the SBR approaches unity, image quality deteriorates rapidly with increasing depth ^13,14^. Therefore, the depth at which the SBR = 1 is typically defined as the depth limit of MPM. In 2-photon microscopy (2PM) with 920 nm excitation, this depth limit typically occurs at a depth of approximately 0.6 to 0.9 mm in vivo (or 4 to 6 EALs), depending on the labeling density. Owing to its higher-order nonlinear excitation and reduced scattering at longer wavelengths, 3-photon microscopy (3PM) enabled greater imaging depth than 2PM. Previous theoretical analysis showed that the depth limit of 3PM is ∼ 8 EALs at close to 100% labeling density, and experiments in tissue phantoms have shown no degradation of the point-spread function (PSF) even at ∼ 10 EALs depth using 3-photon excitation (3PE) at 1300 nm ^16^. For in vivo imaging of the mouse brains, however, the maximum imaging depth achieved by 3PM at 1300 nm is ∼ 1.2 mm for functional and ∼ 1.6 mm for structural imaging ^3,17^, or 5 to 6.5 EALs (accounting for the more scattering white matter layer), which is far from reaching the 3P depth limit. Indeed, the SBRs at the current maximum depths remain high, and it is the signal strength rather than the SBR that limits the achievable imaging depth in 3PM today ^6,12,17,18^.

The choice of excitation wavelength is critical for maximizing imaging depth in brain tissue, with two defined maxima of EALs centered near 1300 nm and 1700 nm. 3PE at 1300 nm is widely used for exciting blue and green fluorophores, including genetically encoded calcium indicators (GECIs) such as GCaMPs ^19^.

Here, we present a 1300-nm 3PM system optimized for maximal imaging depth through a combination of improvements in excitation and collection. The optimum excitation source should provide high pulse energies at the focus for efficient 3PE but still sufficiently below the saturation pulse energy and the nonlinear damage threshold for the imaging depths in the tissue, and average power (which depends on the repetition rate) levels that are tailored to avoid significant sample heating (Supplementary Fig. 1). We designed a 3PM system that delivers ∼1 μJ laser pulses at the brain surface with 100 kHz repetition rate (Supplementary Fig. 2), which is sufficient for imaging GCaMP8s at 2 mm depth. Compared to the typical 3PM using ∼ 0.1 μJ pulse energy and 1 MHz pulse repetition rate at the brain surface, our system achieves ∼ 100x higher signal generation at ∼ 2 mm depth. Furthermore, we used large field-of-view (LFOV) optics to collect the fluorescence signals generated deep in tissue. We repurposed a LFOV 2P/3P microscope ^20^ that has the required high collection numerical aperture across a LFOV. This resulted in a microscope ideally suited to deep-brain 3PM at > 2mm depth with a moderate (∼400 μm) FOV (Supplementary Fig. 2). When compared to a typical 3P microscope, the LFOV microscope achieves greater than tenfold gain in collection efficiency at depths beyond 2 mm. The combined excitation and collection improvement for 3PM at ∼ 2 mm depth is ∼ 1000x when compared to a typical 3P microscope designed for imaging at ∼ 1 mm depth, which enabled us to image brain functions and structures at ∼ 2 mm and beyond.

To demonstrate the performance of this platform, we selected a set of fluorophores with diverse spectra and 3P cross-sections, including Qdot-605, fluorescein-dextran, Texas-Red, and GCaMP8s. We determined that Qdot-605 requires pulse energies ∼ 0.25 nJ at the focus, while fluorescein-dextran, Texas-Red and GCaMP8s require approximately 2 nJ at the focus. Both are below the threshold for nonlinear damage at the focus ^12^. To maintain an average power at the brain surface of ∼100 mW to prevent tissue heating ^12^, the optimum laser repetition rate is ∼100 kHz for deep-brain imaging (Supplementary Fig. 1).

We performed structural imaging of adult mouse (11–12 weeks old wild-type C57BL/6) brain through a cranial window and measured the depth dependence of the SBR using the labeled vasculature and neurons (Fig. 1, Supplementary Fig. 3, Supplementary Videos 1–2). The measured EALs (Supplementary Fig. 3A, 6B) for the mouse brain tissue in our experiments are consistent with previous results ^15^. Using Qdot-605–labeled vasculature, we acquired structural stacks extending to approximately 2.5 mm below the dura. At depths ∼ 2.5 mm (or ∼ 8.2 EALs), the measured SBR approached unity, which indicated that we have reached 3P depth limit for vasculature imaging (Fig. 1, Supplementary Figs. 4–5). This result is consistent with the theoretical prediction for the 3P depth limit of mouse brain vasculature with an excitation NA of 0.5 ^16^. Imaging fluorescein–dextran labeled vascular provided reliable visualization of vasculature to ∼2.2 mm depth (Supplementary Fig. 6, Supplementary Video 3). Furthermore, we confirmed that the resolution in deep tissue is sufficient to resolve a single capillary or neuronal soma (Fig. 1 and Supplementary Fig. 7).

**Fig. 1.**
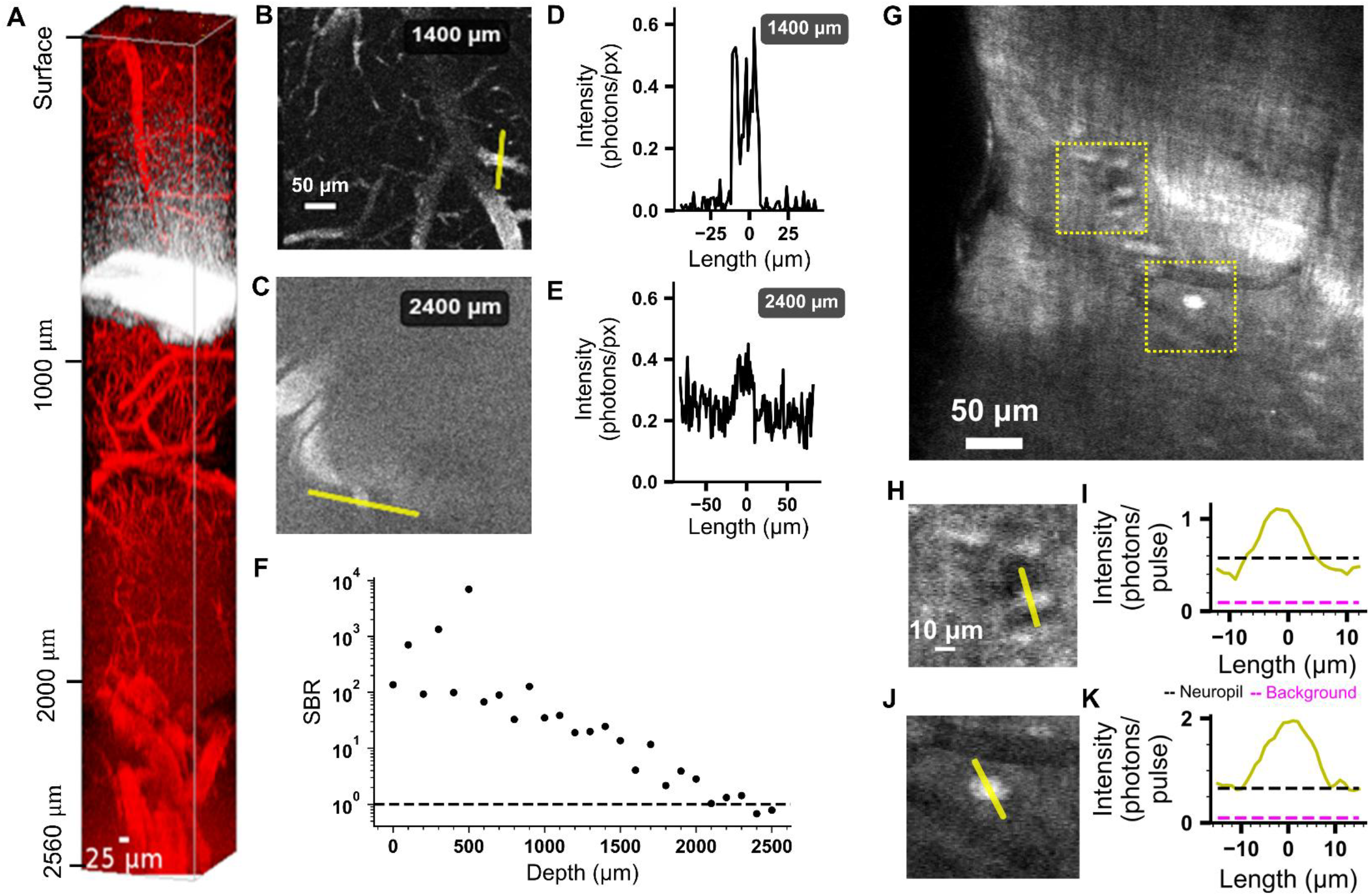
Quantification of SBR for 1300 nm 3PM of mouse brains in vivo. **A**. Imaging Qdot-605-labeled vasculature to approximately 2.5 mm below the pial surface. **B and C**. Representative images of vasculature at a depth of 1400 µm and 2400 µm, respectively, with regions of interest (ROIs) indicated by yellow lines. **D** and **E**. Pixel intensity profiles along the ROIs shown in B and C, respectively, plotted in absolute units of detected photons per pulse. **F**. SBR as a function of depth, calculated by dividing the average signal intensity inside ROIs by the average intensity in adjacent background regions within each slice. **G**. Neural tissue expressing GCaMP8s at a depth of 2000 µm. Distinct somata of individual neurons are visible as bright round structures, whereas blood vessels appear as dark, well-demarcated regions. The boundary of the white matter is visible, curving along the upper right, with neurons apparent at the cingulate cortex–white matter interface. Two regions, highlighted by dotted yellow boxes, are selected for further analysis. **H and J**. ROIs drawn across an individual neuron within the boxed regions in G. **I and K**. Corresponding intensity line profiles across the ROIs in H and J, respectively. Neuropil (black dashed lines) indicated the intensity in pixels adjacent to the neurons and the true background (dashed purple lines) indicated the intensity in pixels within nearby blood vessels.

We performed 3P structural imaging of neuronal architecture in the intact mouse brain beyond 2 mm depth (Fig. 2, Supplementary Figs. 8–11, Supplementary Videos 4–7). Imaging was performed in 8 – 20 weeks old adult male and female transgenic GCaMP8s expressing mice. For structural imaging, volumetric stacks were acquired with a 5 or 10 µm axial step size over a 200 × 200 µm field of view (in some cases larger step size e.g. 50 µm for depth >1800 µm), spanning cortical layers through subcortical hippocampal regions (Fig. 2A–B, Supplementary Figs. 8A–B, 9A–B, Supplementary Videos 4–5). Beneath the cortical layers, hippocampal structures were readily visualized, including CA1 pyramidal neurons (Figs. 2A and 2C, Supplementary Figs. 8A–C and 9A–C) and the trilaminar organization of the dentate gyrus (Fig. 2A, Supplementary Fig. 8B). Cell bodies exhibited characteristic nuclear exclusion of GCaMP8s and well-defined somatic boundaries, enabling identification of pyramidal neuron morphology and local cytoarchitecture (Fig. 2C). The granule cell layer, molecular layer, and polymorphic layer displayed distinct cellular density patterns consistent with known hippocampal anatomy. In the third harmonic generation (THG) channel, myelinated fiber tracts were prominently visible (Fig 2A and B). The external capsule exhibited strong THG contrast with a fibrous morphological characteristic of densely myelinated axonal bundles, providing an intrinsic structural landmark separating cortical and hippocampal regions. Additional imaging in the cingulate cortex (Cg, also referred to as, posterior prefrontal cortex) demonstrated penetration depths up to ∼ 2.3 mm below the dura (Figs. 2F–G, Supplementary Figs. 10–11, Supplementary Videos 6-7). Neuronal somata were clearly visible in medial cortical regions corresponding approximately to secondary motor cortex (M2) and cingulate cortices (shallow-Cg1 and deep-Cg2). The measured SBR was greater than 10 even at 2000 µm depth (Fig. 1G–K). We further verified the EAL with dual color (GCaMP8s + Texas-Red) imaging using a single wavelength at 1332 nm ^21,22^ (Supplementary Fig. 12, Supplementary Video 8). Due to the small amount of residue 2P excitation of Texas Red at 1332 nm, the SBR of Texas Red labeled blood vessel approached unity at ∼ 2000 µm depth ^22^. White-matter tracts became visible laterally at approximately 1790 µm depth and gradually curved medially with increasing imaging depth (Supplementary Fig. 11), consistent with known anatomical trajectories of subcortical projection fibers. The THG channel revealed continuous visualization of these white-matter fibers down to the maximum imaging depth, highlighting the ability of this 3PM system to resolve both neuronal cell bodies and myelinated axonal structures deep within intact brain tissue (Supplementary Fig. 11C). The ability to clearly resolve neuronal morphology, cortical layering, hippocampal cytoarchitecture, and deep white-matter fiber organization beyond 2 mm depth demonstrates that this 3PM approach enables high-resolution structural imaging across cortex, hippocampus, and subcortical white matter in the intact mouse brain.

**Fig. 2.**
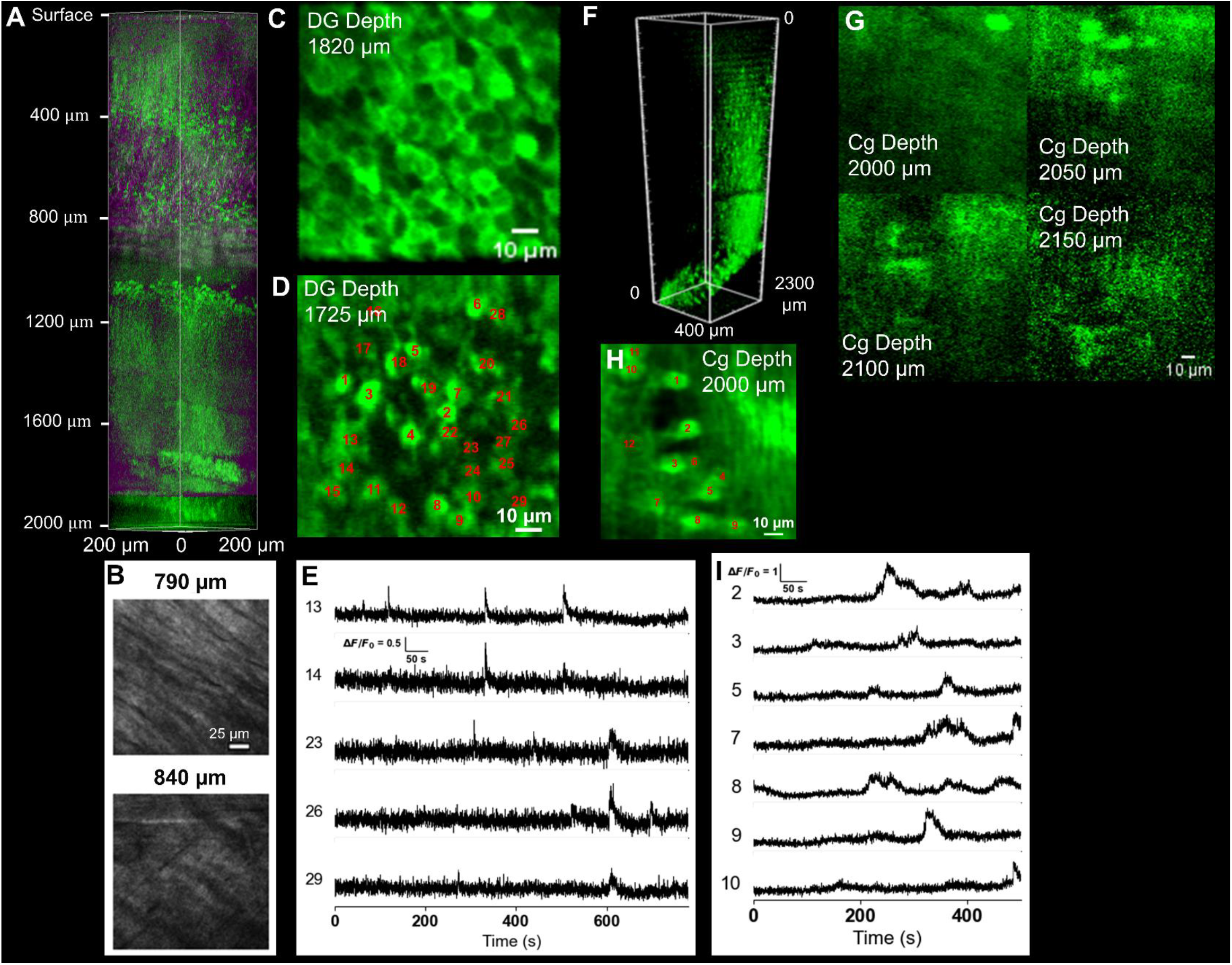
3PM of GCaMP8s transgenic mouse brains. **A**. Three-dimensional reconstruction of in vivo structural imaging in the intact mouse brain, visualizing anatomical features and cytoarchitecture down to 2.0 mm below the pial surface. Distinct cortical layers, as well as the CA1 and dentate gyrus (DG) regions of the hippocampus are visible (green channel). White matter is visible at around ∼800 µm depth (THG channel in magenta). **B**. XY sections at two depths for the THG channel showing the white matter tract. **C**. Single XY section at a depth of 1820 µm within the DG, highlighting the characteristic honeycomb-like arrangement of densely packed granule cells and neural processes. **D**. In vivo functional calcium imaging of DG neurons; identified neuronal soma are annotated at 1725 µm depth. **E**. Representative fluorescence traces from selected neurons within the DG demonstrate spontaneous calcium transients. Additional examples from the same animal and brain region are provided in Supplementary Figures 14-17. Up to 44 neurons could be simultaneously recorded from a 100 µm × 100 µm field of view (120 × 120 pixels) at 6.18 Hz. **F**. 3D reconstruction of a separate imaging session, extending to 2.3 mm depth. **G**. Four XY sections at depths ≥ 2.0 mm. **H**. In vivo calcium imaging; identified neuronal soma are annotated at depth 2.0 mm. **I**. Representative fluorescence traces from selected neurons in H.

We next performed high-resolution functional imaging of neuronal activities in deep brain areas. We recorded spontaneous activity at depths ∼1600 μm (Supplementary Fig. 13, Supplementary Videos 9–10), ∼1725 μm (Figs. 2D–E, Supplementary Figs. 14–16 and 17A–C, Supplementary Videos 11–18) and ∼1820 μm (Supplementary Figs. 17D–F) for deep dentate gyrus (DG) and at 2000 μm (Figs. 2H–I, Supplementary Figs. 17G–I, Supplementary Videos 19–20) for deep cingulate cortex (Cg, also referred to as posterior prefrontal cortex) at frame rates up to 6.18 Hz, imaging populations of up to 44 neurons (Supplementary Fig. 17D) within a 100 × 100 μm FOV (120 x 120 pixels/frame). The measured baseline photon counts (*F*_0_) for all neurons in DG at depth 1725 μm, 1820 μm and Cg at depth 2000 μm substantially exceeded 150 photons/neuron/sec (Supplementary Fig. 17), which indicated high recording fidelity at depth.

We verified that an imaging speed of 6.18 Hz was capable of detecting a single Ca-transient for GCaMP8s labeled neurons in CA1, a region with well-established imaging benchmarks. At ∼1,090 µm depth, reliable Ca-transient detection was achieved with 100 kHz (Supplementary Fig. 18) laser repetition rate which produced *F*_0_ > 150 photons/neuron/sec.

In summary, by systematically optimizing both excitation and collection of 3PM, we extend the imaging depth by > 1.5x relative to previous demonstrations, thus enabling noninvasive, high-resolution observation of structure and function in deep brain regions previously inaccessible by optical methods. This platform will facilitate longitudinal studies of deep circuit dynamics and disease in neuroscience and has broad implications for deep-tissue imaging in other biomedical fields.

## Acknowledgements

We thank members of the Xu research group for their help and constructive discussions, especially Alan Liang and Selin Gildiz for help with animal handling. We also thank Jamien Shea and Nilay Yapici for discussions. M.R. and C.L. were supported by Cornell Neurotech Mong Fellowship. National Institutes of Health U01NS128660.

## Author Contributions

M.R. and C.X. conceived the study. M.R., C.L. and D.G.O built the microscope. M.R. and C.L. performed the experiments, M.R. analyzed the data. M.R. wrote the first draft. M.R., C.L., D.G.O. and C.X. prepared the final manuscript.

## Methods

### Excitation source

The excitation source is an optical parametric amplifier (OPA, Opera-F, Coherent) pumped by a 1,035 nm laser operating at 0.1 to 1 MHz (Monaco, Coherent). The OPA generated femtosecond pulses centered at 1300 to1350 nm. A two-prism (N-SF11 glass) compressor was used to compensate for the normal dispersion of the optics, resulting in a pulse duration of ∼60 fs (measured by second-order interferometric autocorrelation) under the objective. A half-wave plate (AQWP10M-1600, Thorlabs) and a polarization beam splitter (PBS124, Thorlabs) were placed before the pulse compressor to control the excitation power.

### Imaging system

Efficient fluorescence collection from deep within scattering tissue remains a major challenge, as existing microscopes exhibit low collection efficiency beyond ∼1 mm depth. The collection efficiency scales approximately with NA^2^·FOV^2^ (with respect to signal collection), while scattered fluorescence exits the tissue over a diffuse region whose FWHM is ∼1.5× the imaging depth ^13^. Thus, at 2 mm depth the emission spreads over ∼3 mm diameter, requiring a collection FOV of ∼3.3 mm for a 0.3 mm scanned region. However, commonly used objectives (e.g., the Olympus 25×, 1.05 NA objective) provide only ∼1 mm collection FOV (80% throughput), leading to ∼11-fold reduced fluorescence collection efficiency at 2 mm depth. To improve collection at depth, we took advantage of a recently developed large field-of-view (LFOV) 2P/3P microscope ^20^ that has the required high collection numerical aperture (NA ∼ 1.0) across a LFOV (∼ 4 mm at 80% throughput).

Raster scanning was achieved using a 5 mm-aperture non-conjugated galvo-galvo scanner (Saturn 9B, ScannerMax). Custom scan lens (f = 60 mm) and tube lens (f = 200 mm) (Special Optics, Navitar) relay the beam to the objective back pupil (f = 15 mm, Special Optics, Navitar).The LFOV microscope had a water-immersion objective (maximum excitation NA 0.75, collection NA 1.0) that was underfilled to an effective excitation NA of ∼0.55 for three-photon excitation.

Fluorescence and THG signals were epi-collected through the objective and reflected by a primary dichroic (FF705-Di01-77x108, Semrock) into the detection arm. Signals were spectrally separated using a 470-nm dichroic (FF470-Di01-77x108, Semrock); green fluorescence (e.g., GCaMP8s) and THG was filtered by green (FF01-520/70, Semrock) and blue (FF02-435/40, Semrock) filters, respectively, and detected with two GaAsP PMT (H15460-40, Hamamatsu). For Texas Red and Qdot-605 imaging, the red fluorescence signal was separated using a 580-nm dichroic (FF580-FDi01-77x108, Semrock) and filtered by a red filter (FF01-607/70, Semrock). PMT currents were converted to voltage signals by a transimpedance amplifier (HCA-200 M-20 K-C, Femto), and digitized with vDAQ board (Vidiro). The signal was low pass filtered with vDAQ built-in anti-aliasing 40MHz filter. Signal gating was performed with 24 ns sampling window (4 sampling points per pulse at 125 MHz sampling rate) synchronized with laser clock signal from the Monaco laser. Control and acquisition were performed using ScanImage (Vidrio) in MATLAB (MathWorks), with sample positioning provided by a motorized stage (MP-285, Sutter). The system achieved shot-noise-limited performance.

### Optimization of the pulse energy at the focus and the laser repetition rate

3P fluorescence signal can be quantified according to the following set of equations where ⟨*F*(*t*)⟩ is the time-averaged 3P fluorescence signal, *τ* is the pulse width, *f* is the laser repetition rate, *P*_focus_ is the average power at the focus, *P*_surface_ is the average power at the brain surface, *E*_focus_ is the pulse energy at the focus, *z* is the imaging depth, and EAL is effective attenuation length of the brain tissue.

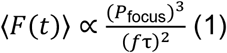

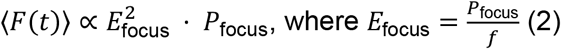

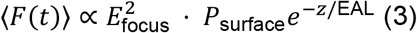

At *λ* = 1300 nm, the maximum three-photon signal is achieved by maximizing *P*_surface_ and *E*_focus_, which are limited, respectively, by brain heating and fluorophore saturation or nonlinear damage, and typical maximum values using a high-NA objective lens are: *P*_surface_ ∼ _*1*00_ mW, *and E*_focus_ ∼ _2_ nJ.

Thus, the pulse energy at the brain surface as a function of imaging depth *z* is given by:

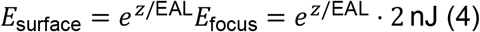

The corresponding repetition rate *f* at depth *z* follows:

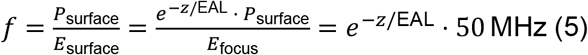

### Depth calibration

Imaging depths were recorded as the raw axial displacement of the objective lens. Due to the higher refractive index of brain tissue relative to water, the reported depths slightly underestimate actual tissue depths by approximately 5–10% ^12^.

### Animal procedures

All animal procedures were conducted in accordance with protocols approved by the Cornell University Institutional Animal Care and Use Committee. Wild-type C57BL/6J male mice (11–12 weeks old, The Jackson Laboratory) were used for blood vessel imaging following retro-orbital injection of Qdot-605, or fluorescein (dextran conjugate, 70 kDa, Invitrogen).

Neuronal imaging was performed in transgenic CamkIIa-GCaMP8s mice (8–20 weeks old). Transgenic mice were generated by crossing B6;129S-Igs7tm2(tetO-GCaMP8s,CAG-tTA2)Genie/J (JAX stock #037719; RRID:IMSR_JAX:037719), commonly known as TIGRE2-jGCaMP8s-IRES-tTA2, with B6.Cg-Tg(Camk2a-cre)T29-1Stl/J (JAX stock #005359; RRID:IMSR_JAX:005359), commonly referred to as T29-1. Both strains were obtained from The Jackson Laboratory. This was also used for dual color imaging following a retro-orbital injection of Texas-Red (dextran conjugate, 70 kDa, Invitrogen).

### Surgical preparation

Cranial windows, 5-8 mm in diameter were implanted for in vivo imaging. The window was centered at 2.5 mm lateral and 2 mm caudal to bregma for imaging the DG area and was centered at 0 mm lateral and -0.5 mm caudal to bregma for imaging the CG area. Anesthesia was induced with 3-5% isoflurane and maintained at 1– 2% during surgery. Body temperature was held at 37.5°C using a feedback-controlled heating blanket (Harvard Apparatus), and ophthalmic ointment was applied to prevent corneal drying. Pre-operative dexamethasone (0.2 mg/kg, subcutaneous) and bupivacaine (local, subcutaneous) were administered, with ketoprofen (5 mg/kg) provided post-operatively and for two subsequent days. The cranial window was sealed with a circular coverslip fixed in place with dental acrylic. Imaging was initiated immediately post-surgery. No statistical methods were employed to predetermine sample size, and randomization or investigator blinding was not utilized.

### Imaging protocol

During imaging, animals were maintained under light anesthesia with 1–1.5% isoflurane in oxygen, titrated to sustain a respiratory rate of ∼2 Hz, and body temperature was maintained at 37.5°C. Animals received ophthalmic ointment, and were positioned on a three-axis motorized stage for precise navigation under the microscope.

Three-photon structural imaging stacks were acquired at 400 × 400 pixels. For blood vessel signal-to-background ratio (SBR) measurements, fluorescein, Qdot 605, or Texas Red was administered via retro-orbital injection prior to imaging. Qdot 605 stock solution was injected at a volume of 120 µL. Fluorescein (10–12 mg) was dissolved in 120 µL saline before injection. Texas Red (10 mg) was dissolved in 120 µL saline and injected at the same volume.

Three-photon imaging of neural activity was conducted at a frame size of 100 µm x100 µm (120 × 120 pixels) and a frame rate of 6.18 Hz, with imaging sessions lasting up to 60 minutes. We performed multiple sessions in the same animal lasting up to 5 hours with 0–5 mins interval between sessions.

Similar to previous reports ^19^, GCaMP8s exhibits some photobleaching during the beginning of the imaging session which reaches a steady state after the first 10-15 mins, and hence the activities presented here are taken after the first 15 mins of recording.

All imaging parameters have been summarized in Supplementary Table 1.

### Photon counting

ROIs were manually identified in Fiji. Absolute photon count rates were determined by multiplying integrated pixel values by a calibration factor, derived from parallel measurements of photon count rate (SR400, Stanford Research Systems) and image pixel values on a standard fluorescein sample. Calibration was performed by parking the excitation focus on a fluorescein solution and sequentially conducting photon counting and standard imaging measurements. Linearity between pixel intensity and photon counts was validated by varying the excitation laser power. The ratio of photon counts to pixel values under linear conditions was defined as the conversion factor.

Neuronal activity traces were extracted by integrating fluorescence over individual somata (∼100 pixels at the reported field of view).

### Processing of structural images

For structural images, each raw data frame was normalized using histogram stretching, allowing 0.2–0.5% pixel saturation, prior to three-dimensional visualization. Image stacks were rendered using Imaris (Bitplane) or Fiji Volume Viewer after processing with custom Python scripts.

### Processing for activity traces

Prior to analysis, imaging data were corrected for lateral motion using NoRMCorre (https://github.com/flatironinstitute/NoRMCorre). Regions of interest (ROIs) corresponding to individual neurons were manually identified in Fiji and their activity traces were extracted for processing using custom Python script. Fluorescence intensity traces were low-pass filtered with a hamming window of a time constant of 0.29 s. Baseline of the traces (*F*_0_) is defined as approximately the lower 20% of each trace during the whole recording time. Intensity traces (*F*) are normalized according to the formula (*F* − *F*_0_)/ *F*_0_.

Activity videos are presented in supplementary materials either as raw registered videos or filtered videos. The filtering was achieved by a Median 3D filter with X radius = 1.0, Y radius = 1.0 and T radius = 18.0 (∼3 s). First 9 frames and last 9 frames were removed from the filtered videos to remove the edge effect.

### Quantifying d’ for Calcium activity imaging

Assuming exponential decay of calcium transients, calcium transient detectability can be quantified using the discriminability index, *d*^′23,24^.

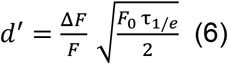

For GCaMP8s, t_1_/_2_ ≈ 200 ms (in vivo), then *τ*_*1/e*_ ≈ 290 ms.

Taking typical published values for jGCaMP8s:

- Δ*F*_*/*_*F* ≈ 0.7(single-AP)
- *τ*_*1/e*_ ≈ _0_._2_9s

| **d'** | **Confidence interval** | **Required $F_0$ (photons/neuron/s)** |
| --- | --- | --- |
| 4.65 | 99% | ≈ **305** |
| 3.76 | 97% | ≈ **199** |
| 3.29 | 95% | ≈ **152** |
| 2.95 | 93% | ≈ **122** |

