## Supplementary materials for "Three photon microscopy of mouse brain structure and function at 2 mm depth and beyond"

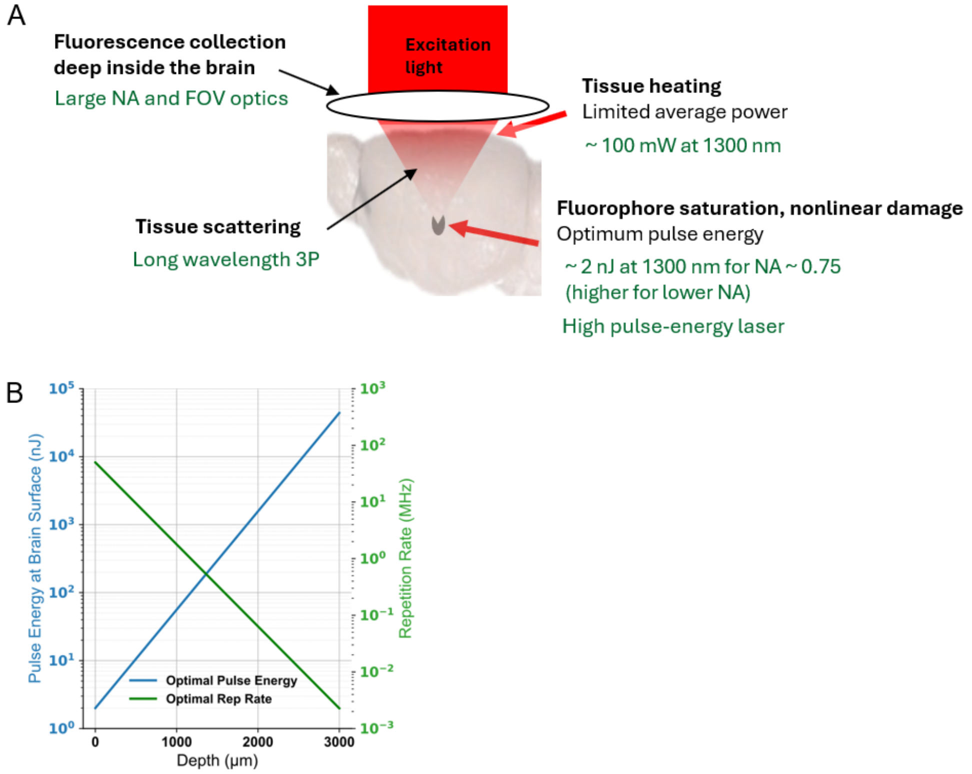


### Supplementary Figure 1

Optimum excitation parameters and microscopy setup for maximum 3P fluorescence signal deep within brain tissue.

A. An illustration of the challenges of deep tissue imaging and the technological advancements required to overcome those challenges.

B. Calculated optimum pulse energy at the brain surface (solid blue line) and pulse repetition rate (dashed green lines) for 3PM at 1300 nm as a function of imaging depth (assuming EAL = 300 μm). The calculations use Eqs. (4) and (5) (see Methods for equations).

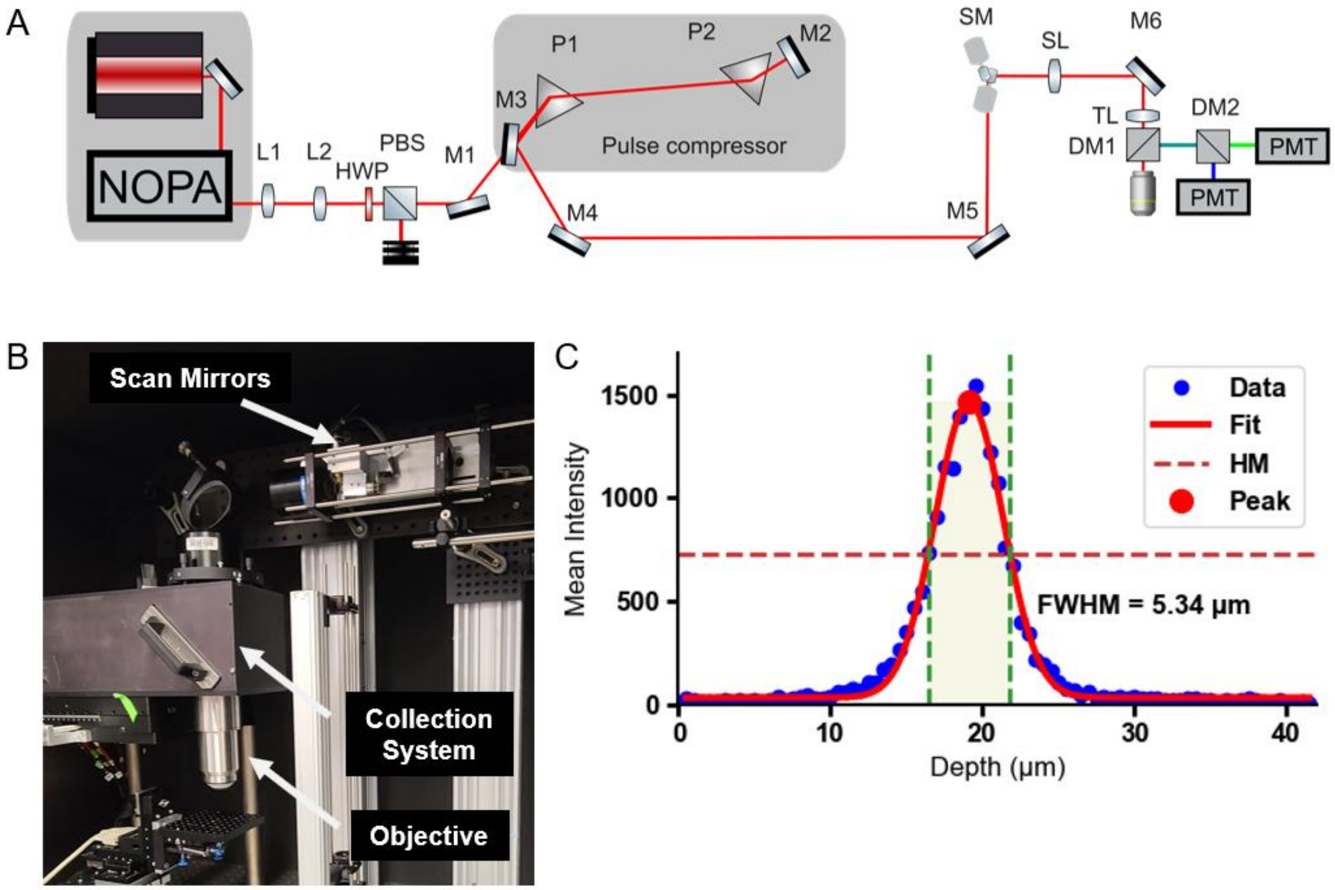


**Supplementary Figure 2**

Experimental setup

A. Schematic of the excitation source, optical path and the collection system. NOPA – non-collinear optical amplifier, L – lens, HWP – half wave plate, PBS – polarizing beam splitter, M – mirror, S – shutter, SM – scanning mirrors, SL – scan lens, TL – tube lens, DM – dichroic mirror, PMT – photomultiplier tube.

B. A photo of the microscope and the imaging stage.

C. The axial resolution of the microscope measured by scanning a thin fluorescent film across the focus. HM – half maximum, FWHM – full width half maximum

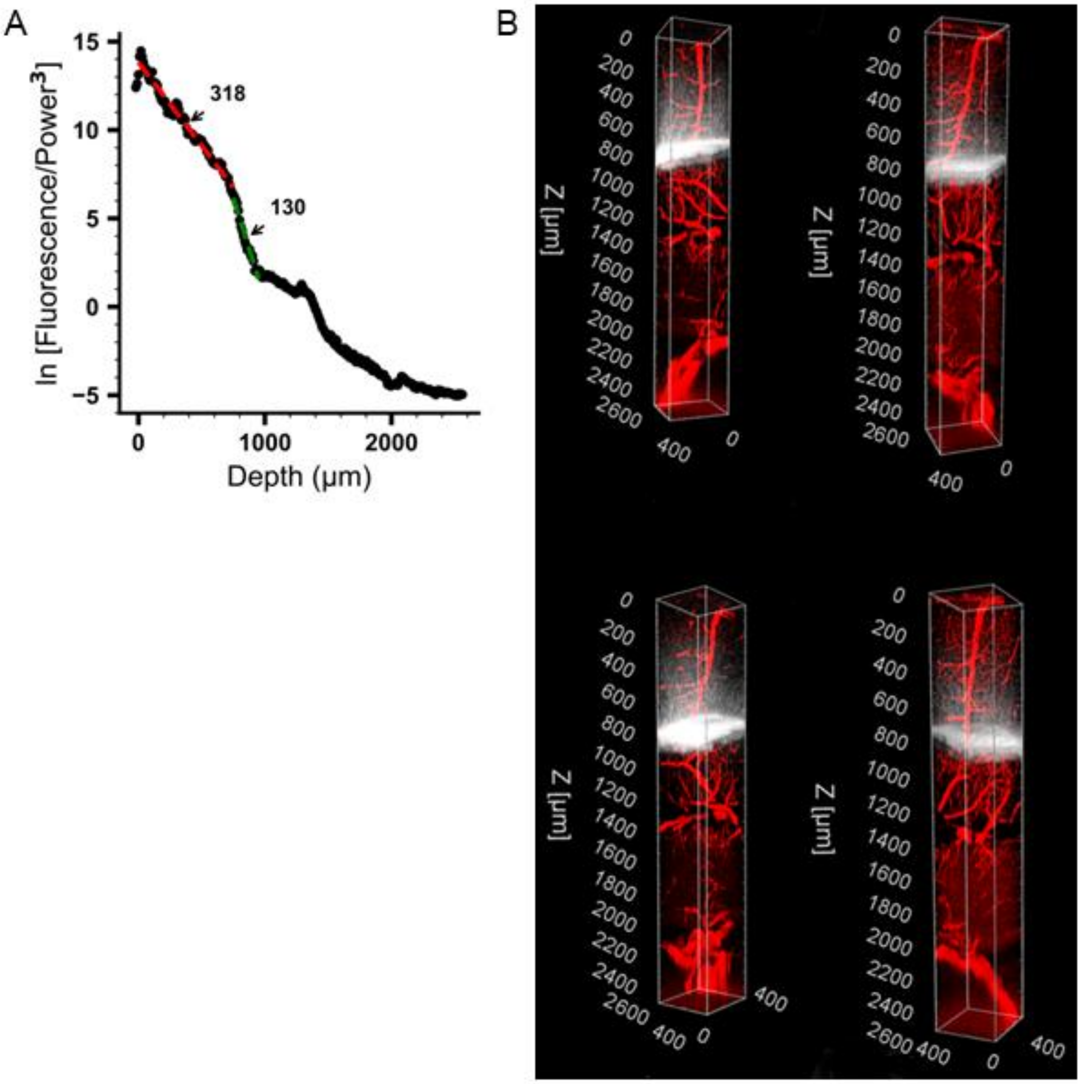


**Supplementary Figure 3**

In vivo 3PM of Qdot-605-labeled vasculature with imaging depths up to approximately 2.5 mm below the pial surface. Brain vasculature was visualized with retro-orbital injection of Qdot-605 and a cranial window placed near bregma AP – 2.5 mm and ML + 2.5 mm.

A. Measuring the effective attenuation length (EAL) by plotting the fluorescence intensity, calculated for the top 0.1% of pixel intensities from each optical section, as a function of imaging depth.

B. Four perspectives of the 3D reconstruction of Qdot-labelled blood vessels.

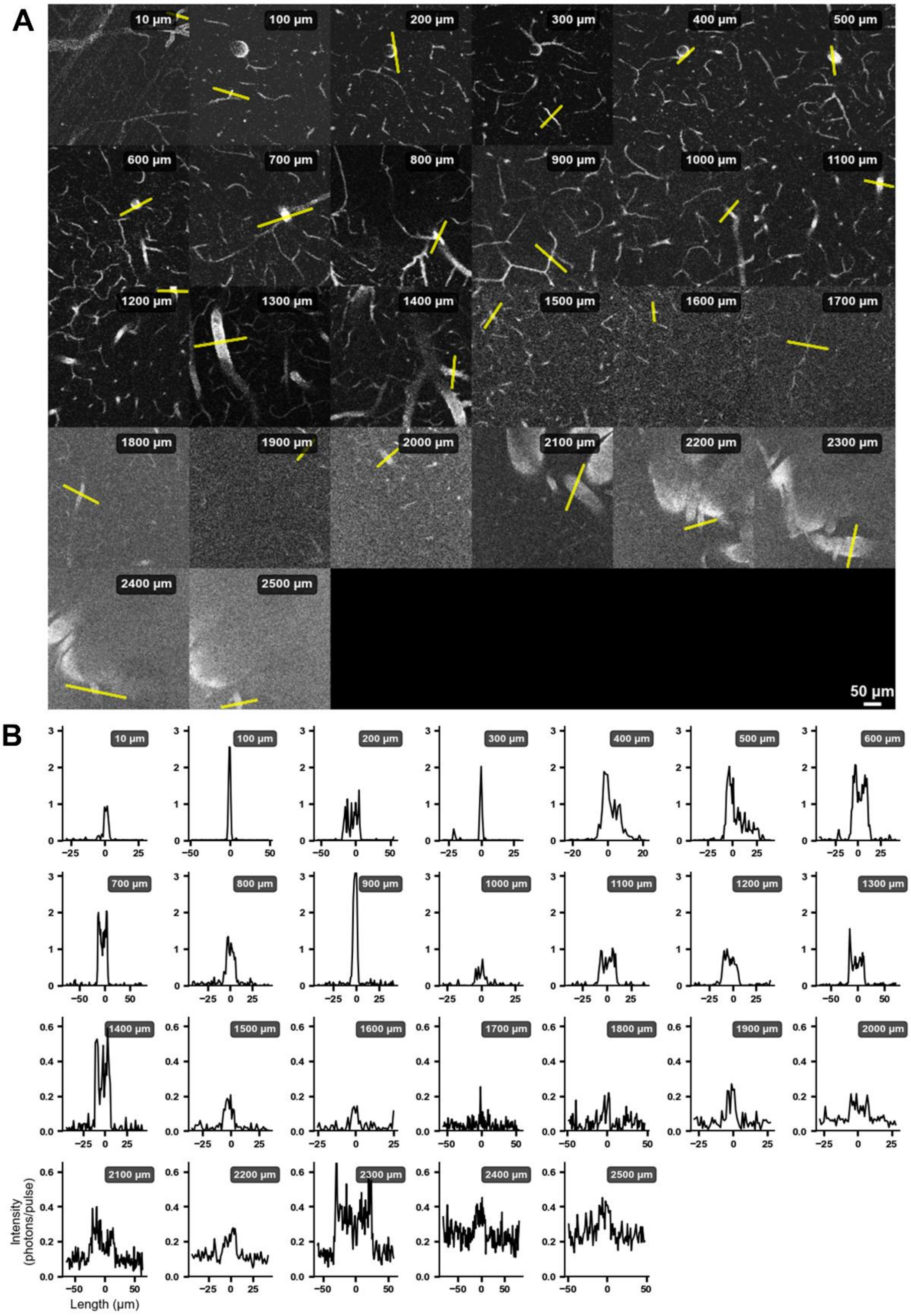


**Supplementary Figure 4**

Quantification of signal-to-background ratio (SBR) for Qdot-605-labelled blood vessels.

A. XY sections of Qdot-605 labelled blood vessels at various depths. ROIs are marked with yellow lines.

B. Line intensity profiles of the selected ROIs.

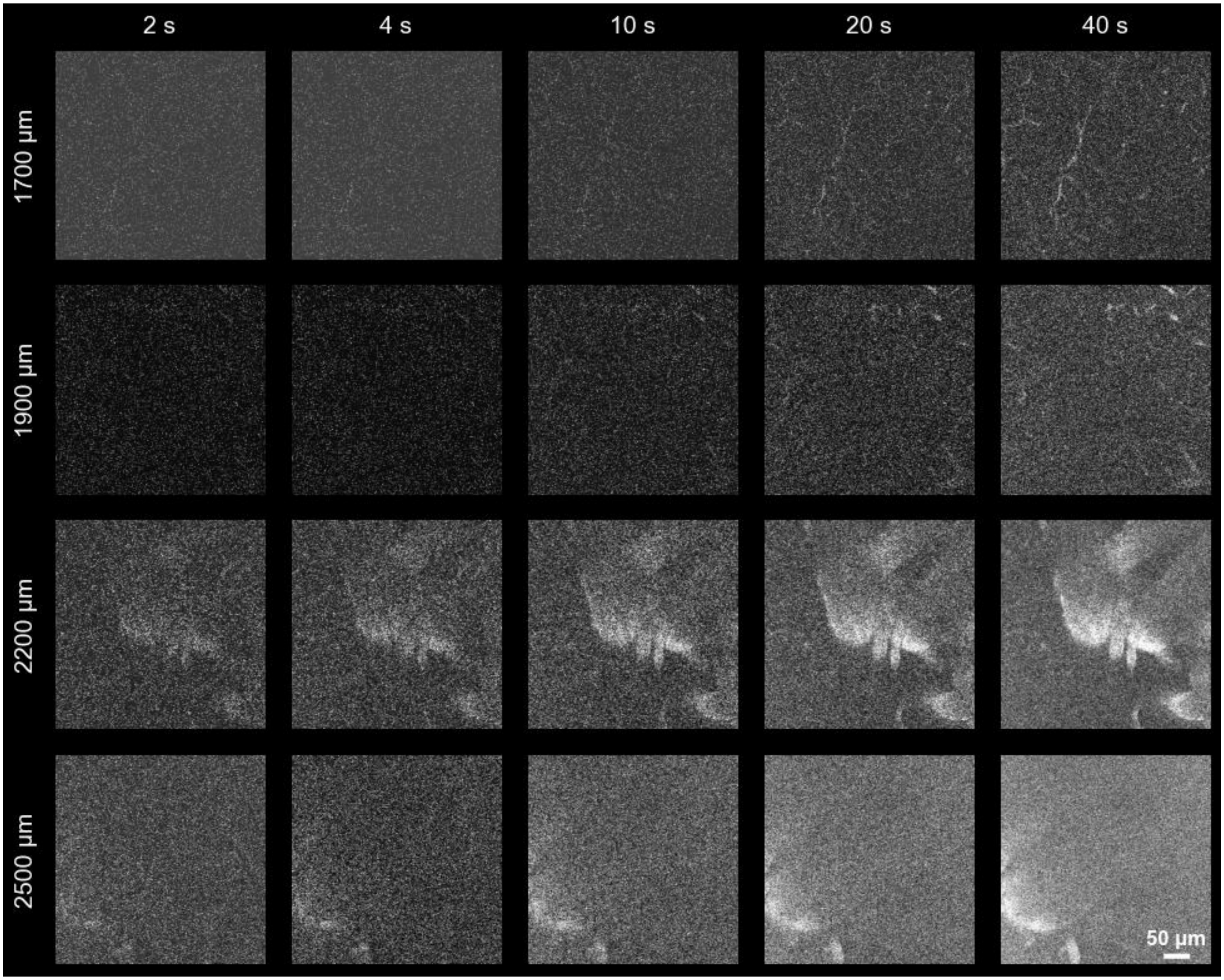


**Supplementary Figure 5**

The impact of frame averaging on deep blood vessel imaging with Qdot-605.

The rows are for different imaging depth (indicated on the left), and the columns are for the images obtained after averaging for the durations indicated at the top. For each frame, raw data were normalized using histogram stretching, allowing saturation of 0.2–0.5% of the pixels.

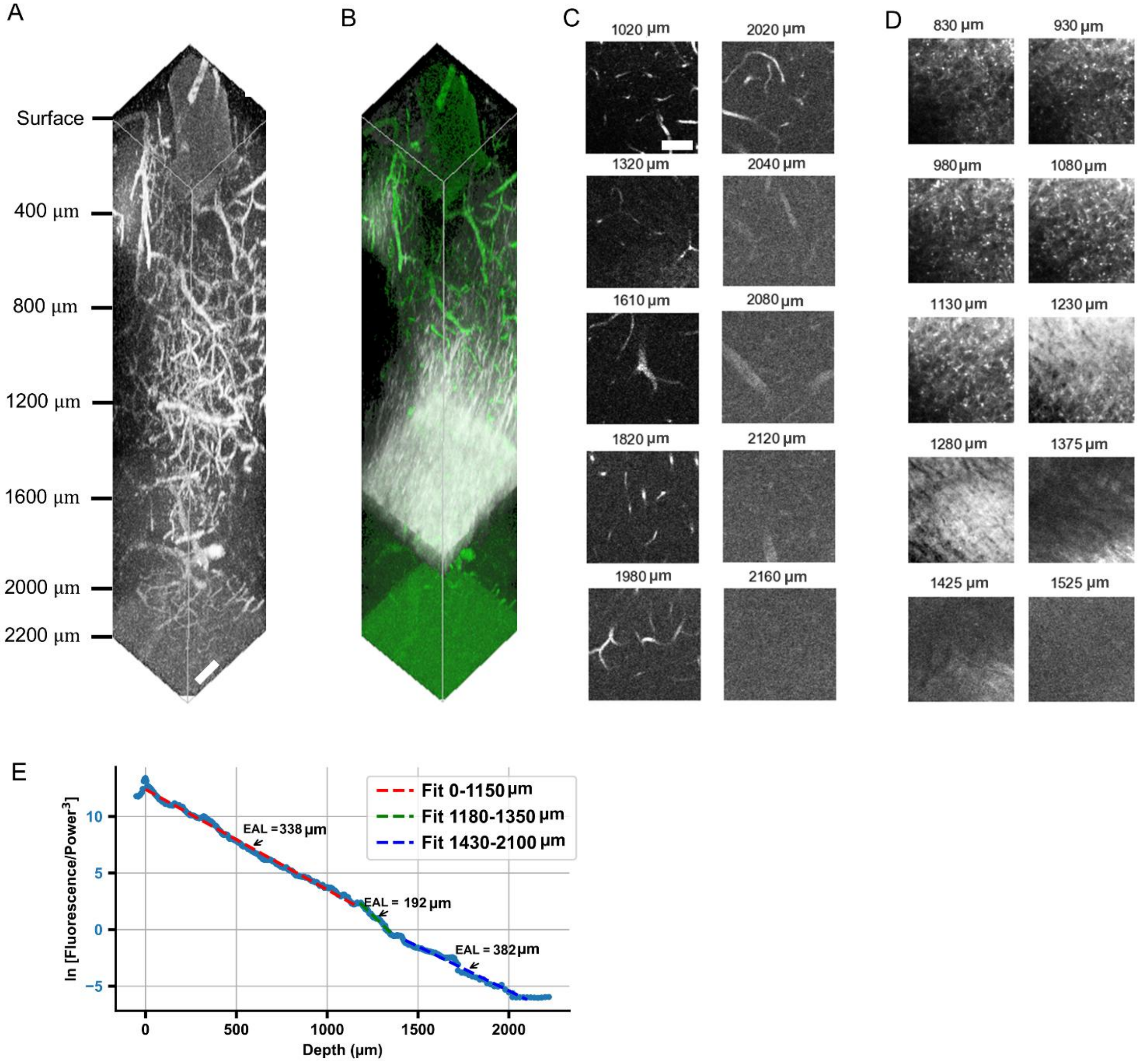


**Supplementary Figure 6**

3PM of fluorescein-labeled blood vessels down to 2.2 mm depth. Each raw data frame was normalized using histogram stretching, allowing saturation of 0.2–0.5% of the pixels.

A. 3P imaging in the mouse brain using fluorescein-dextran retro-orbital injection and a cranial window placed near bregma AP – 1.5 mm and ML + 2 mm.

B. White matter is visualized using the third harmonic generation (THG) channel shown in gray here alongside fluorescein-dextran in green.

C. XY sections of fluorescein-dextran labelled blood vessels at various depths.
D. XY sections of THG channel at various depths.

E. Fluorescence intensity (top 0.1% pixels) as a function of imaging depth in a semi-log plot. The EALs are determined by the slope of the linear fit to the fluorescence intensity data.

All scale bars are 50 µm.

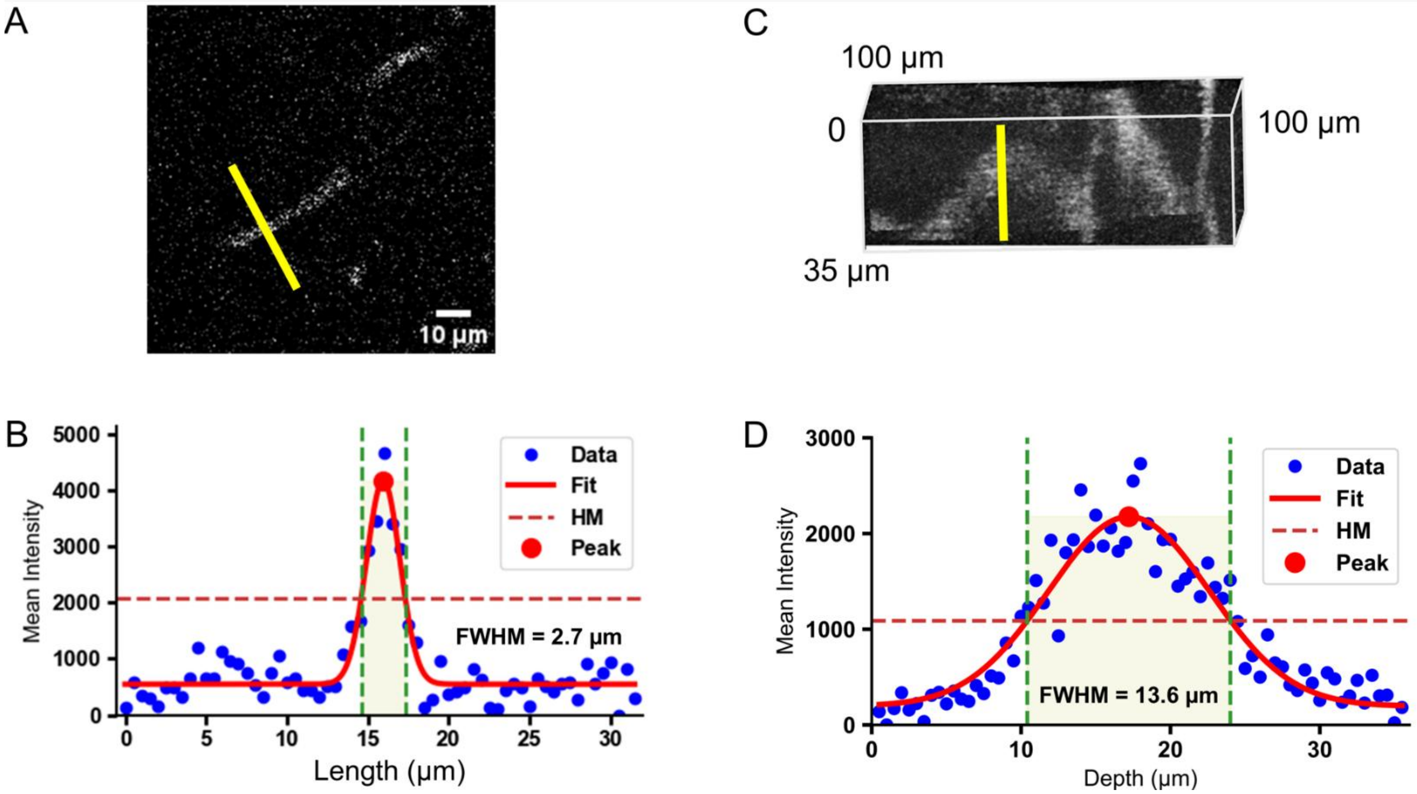


**Supplementary Figure 7**

Characterization of the spatial resolution at ~ 1780 µm depth for a thin capillary labelled with fluorescein-dextran. Taking into account that the minimum diameter of the mouse capillary is 2 to 3 µm, these measurements indicate that the spatial resolution (FWHM) is ~ 1 µm or below laterally and ~ 13 µm axially.

A. A XY section at 1780 µm depth. Yellow line shows the ROI.

B. Lateral line profile for the ROI shown in A.

C. 3D reconstruction of the stack between 1760 and 1795 µm. Yellow line shows the ROI along the axial direction of the same vessel in A. 100 µm, 100 µm and 35 µm represent the extent of the volume in X, Y and Z directions, respectively.

D. Axial intensity profile along the yellow line shown in C.

FWHM – full width half maximum, HM – half maximum

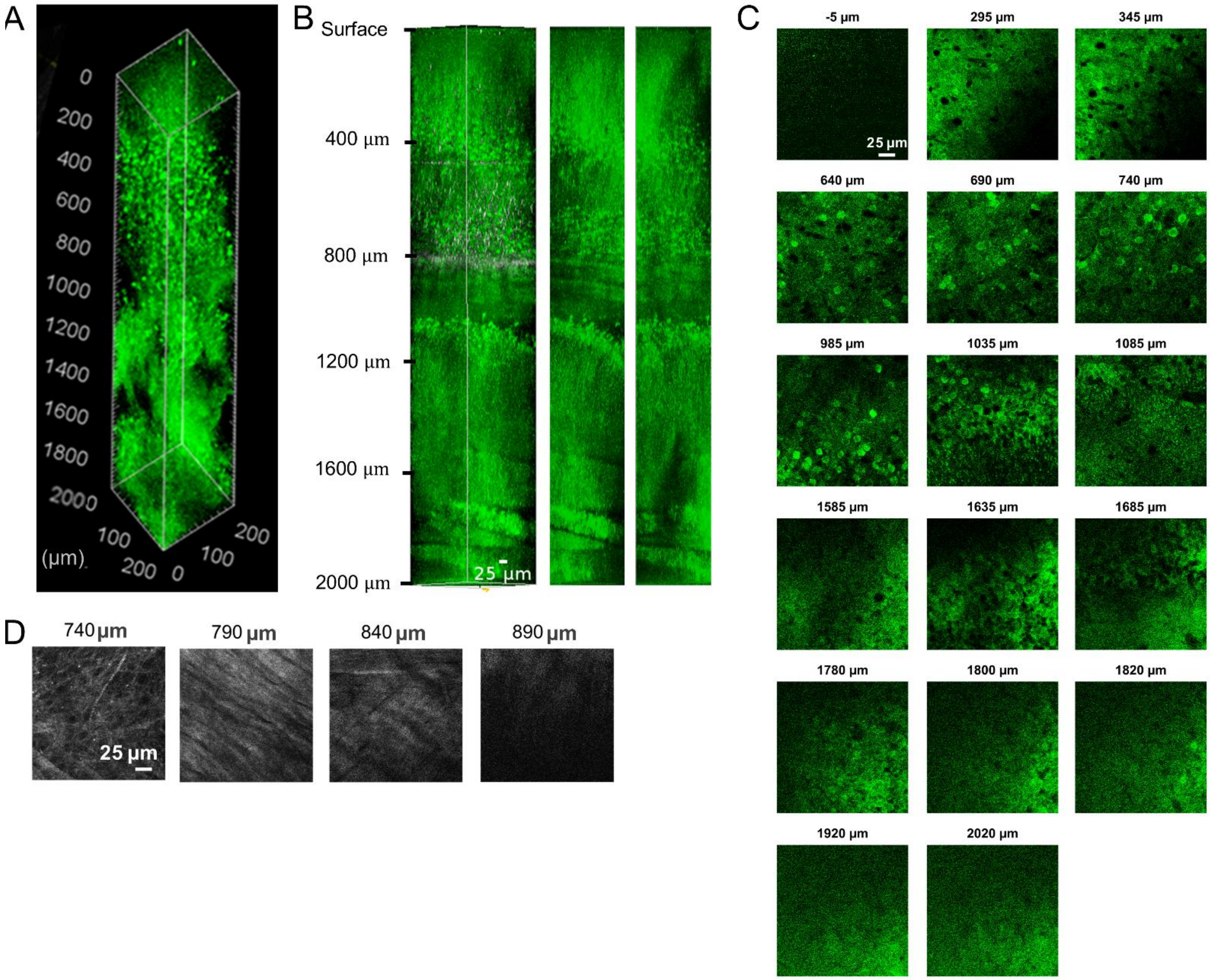


**Supplementary Figure 8**

Anatomical and cellular features across depth for structural imaging of GCaMP8s transgenic mouse in DG area of the hippocampus – mouse #1. Each raw data frame was normalized using histogram stretching, allowing saturation of 0.2–0.5% of the pixels.

A. A 3D reconstruction showing distinct cellular layers from the cortex to the subcortical layers down to the DG area.

B. Layers 2/3, 5, 6, CA1 and trilaminar structure of DG are visible in the 3D reconstruction (left), XZ (middle) and YZ (right) projections.

C. XY sections at various depths for the green fluorescence channel.

D. XY sections at various depths for the THG channel.

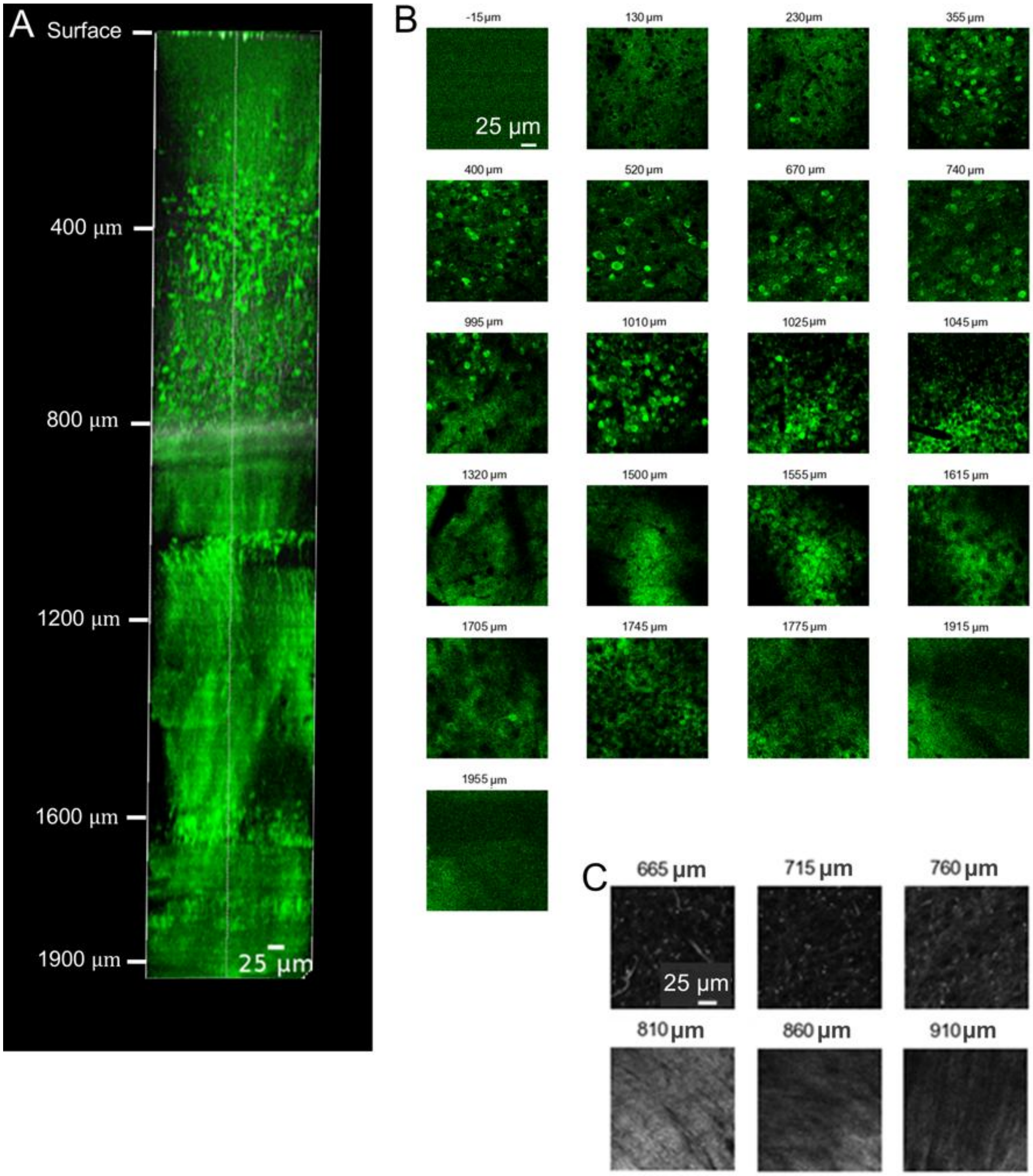


**Supplementary Figure 9**

Anatomical and cellular features across depth for structural imaging of GCaMP8s transgenic mouse in DG area of the hippocampus – mouse #2. Each raw data frame was normalized using histogram stretching, allowing saturation of 0.2–0.5% of the pixels.

A. 3D reconstruction for an imaging stack showing cortical and subcortical cell layers. Layers 2/3, 5, 6, CA1 and trilaminar structure of DG are visible in the 3D reconstruction. It also shows clear distribution of the soma of neurons.

B. XY sections at various depths for the green fluorescence channel.

C. XY sections at various depths for the THG channel.

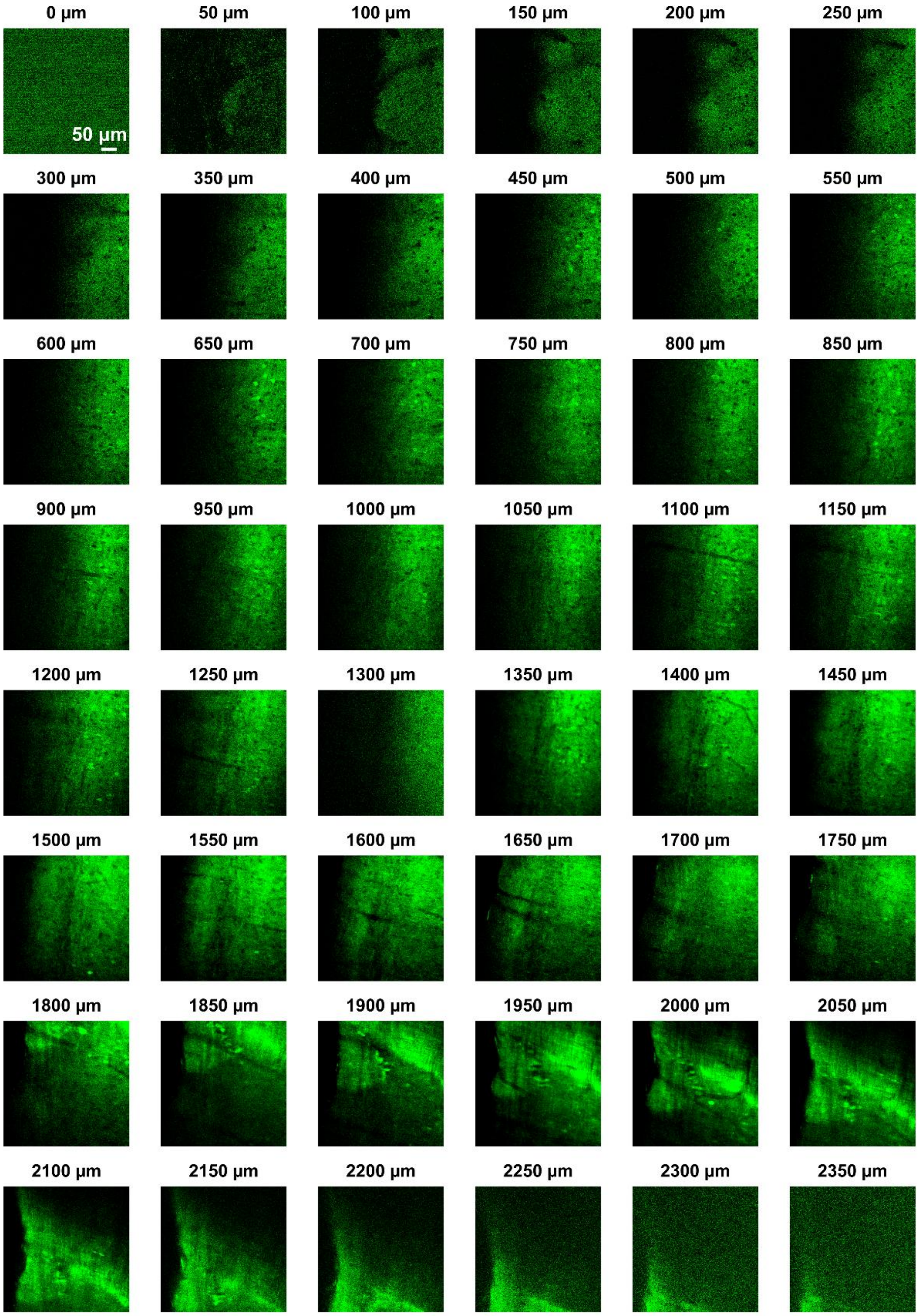


**Supplementary Figure 10**

Anatomical and cellular features across depth for structural imaging of GCaMP8s transgenic mouse in Cg area – mouse #3.

XY sections for the green fluorescence (GCaMP8s) channel at various depths for Fig. 2F to ~ 2.3 mm below the pial surface.

Each raw data frame was normalized using histogram stretching, allowing saturation of 0.2–0.5% of the pixels.

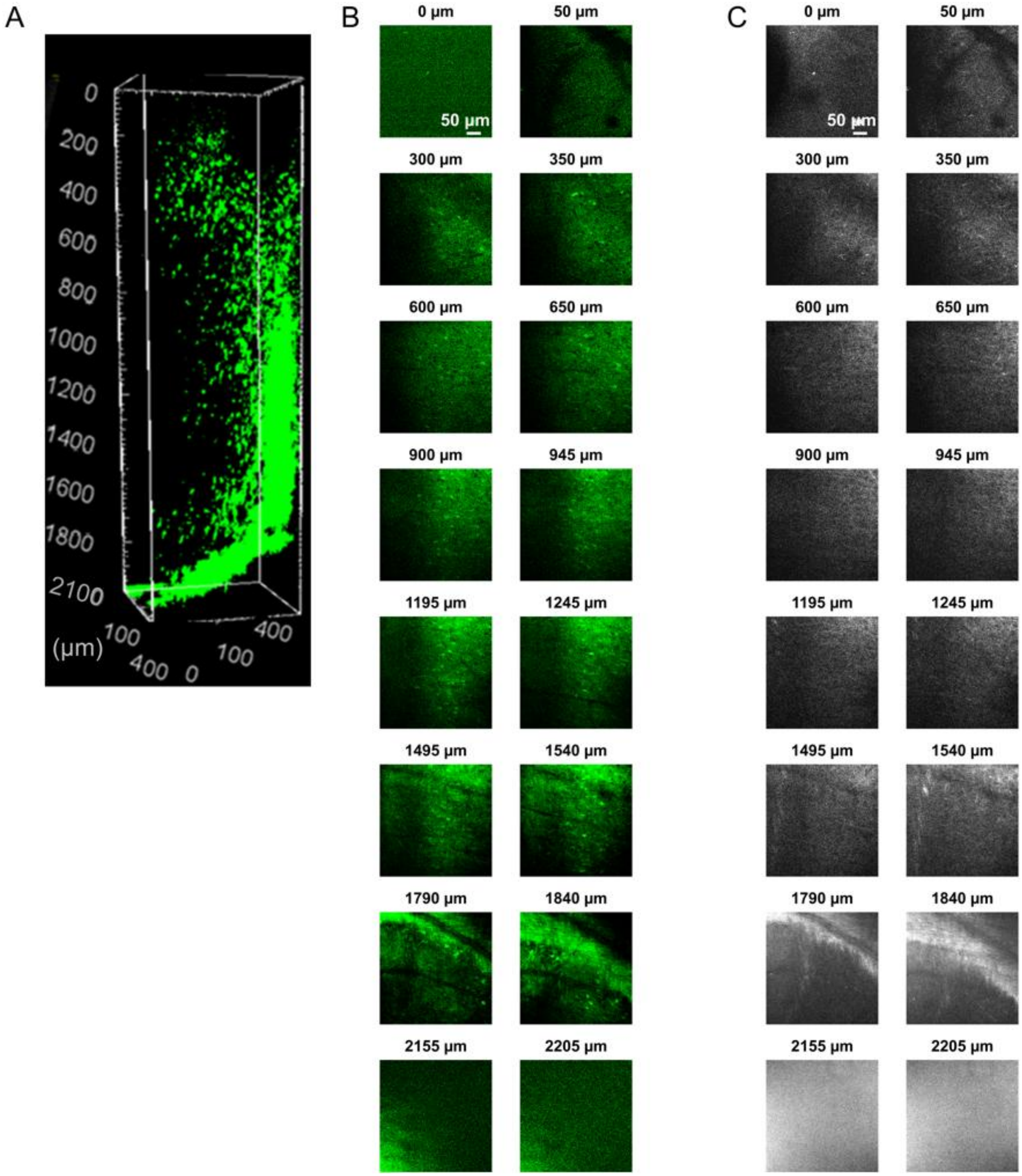


**Supplementary Figure 11**

Anatomical and cellular features across depth for structural imaging of GCaMP8s transgenic mouse in Cg area – mouse #4. Each raw data frame was normalized using histogram stretching, allowing saturation of 0.2–0.5% of the pixels.

A. Structural 3D reconstruction to ~ 2.1 mm depth in cortex. Neurons are apparent as bright, circular soma, while the underlying white matter is distinguished by its dense, fibrous appearance curving in toward the bottom of the image.

B. XY sections at various depths for the green fluorescence channel.

C. XY sections at various depths for the THG channel.

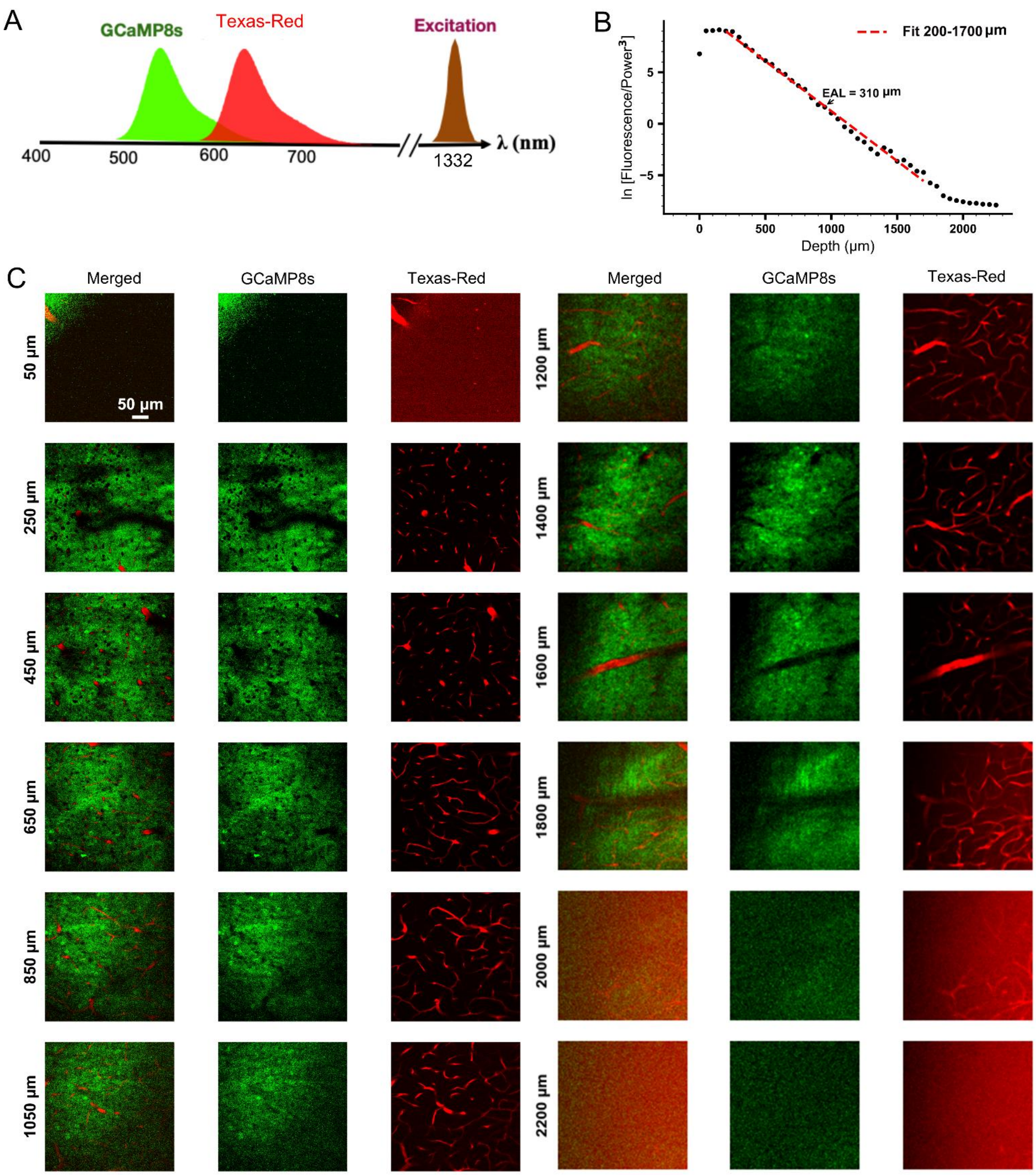


**Supplementary Figure 12**

Dual color imaging at depth with 1332 nm excitation wavelength – mouse #4.

A. A schematic of the approach to excite both a green (GCaMP8s) and red (Texas Red) dye with a single excitation wavelength at 1332 nm.

B. Texas Red fluorescence intensity as a function of imaging depth. Top 0.1% pixels (based on intensity values) were selected for the calculation. The EAL was indicated in the plot.

C. XY sections at various depths for the green (GCaMP8s) and red (Texas Red) fluorescence channel. Left - the combined red and green fluorescence channel, middle - green fluorescence channel, and right - red fluorescence channel.

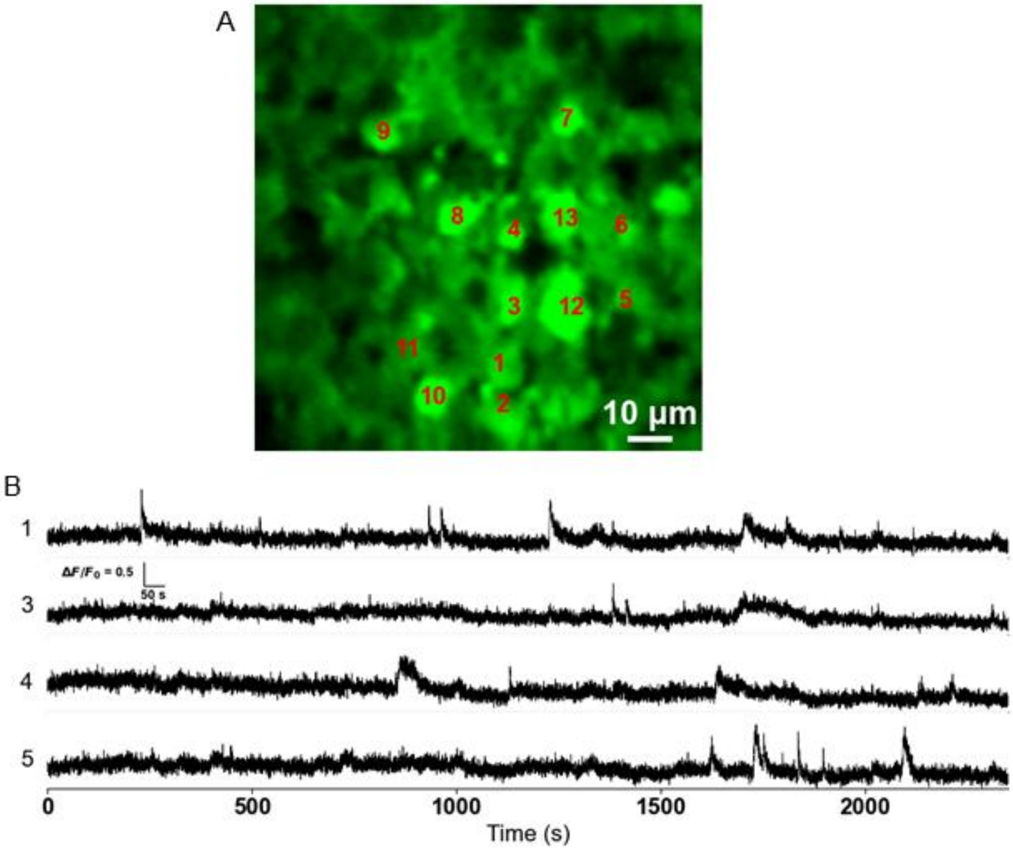


**Supplementary Figure 13**

3P imaging of spontaneous activity in GCaMP8s-labelled neurons in DG area of the hippocampus at depth 1600 µm below the pial surface.

A. Neuronal soma during functional calcium imaging.

B. Fluorescence traces from the active neurons within the DG.

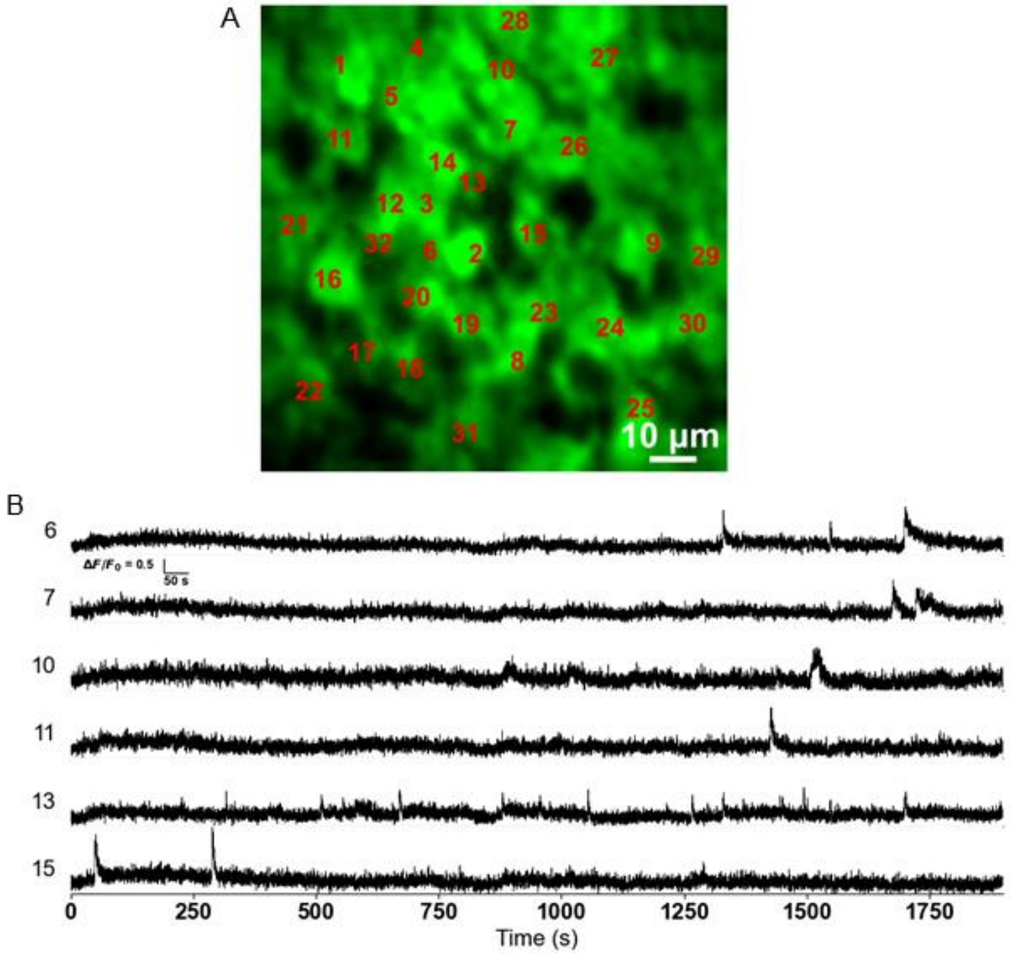


**Supplementary Figure 14**

3P imaging of spontaneous activity in GCaMP8s-labelled neurons in DG area of the hippocampus at depth 1725 µm below the pial surface – session #1 (~1 hr long) from the same FOV as in Fig. 2F (Fig. 2F is session #3).

A. Neuronal soma during functional calcium imaging.

B. Fluorescence traces from the active neurons within the DG.

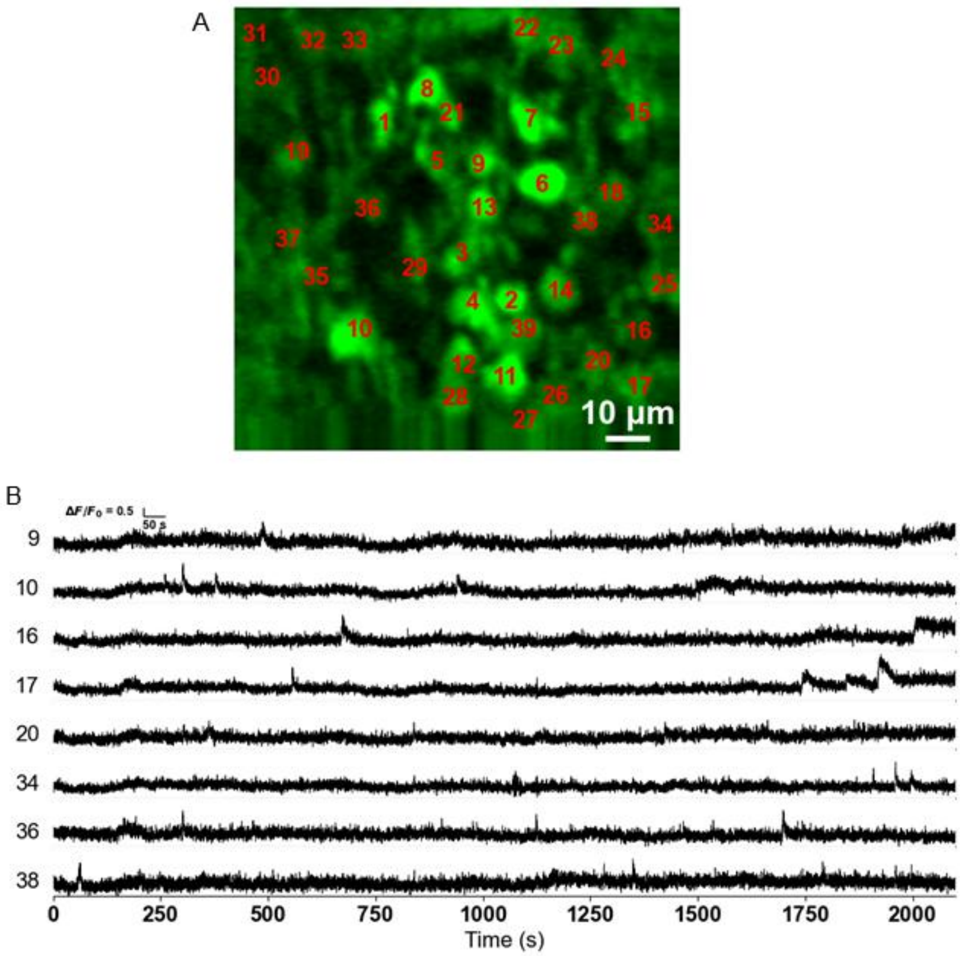


**Supplementary Figure 15**

3P imaging of spontaneous activity in GCaMP8s-labelled neurons in DG area of the hippocampus at depth 1725 μm below the pial surface - session #2 from the same FOV as in Fig. 2F and Supplementary Fig. 14.

A. Neuronal soma during functional calcium imaging.

B. Fluorescence traces from the active neurons within the DG.

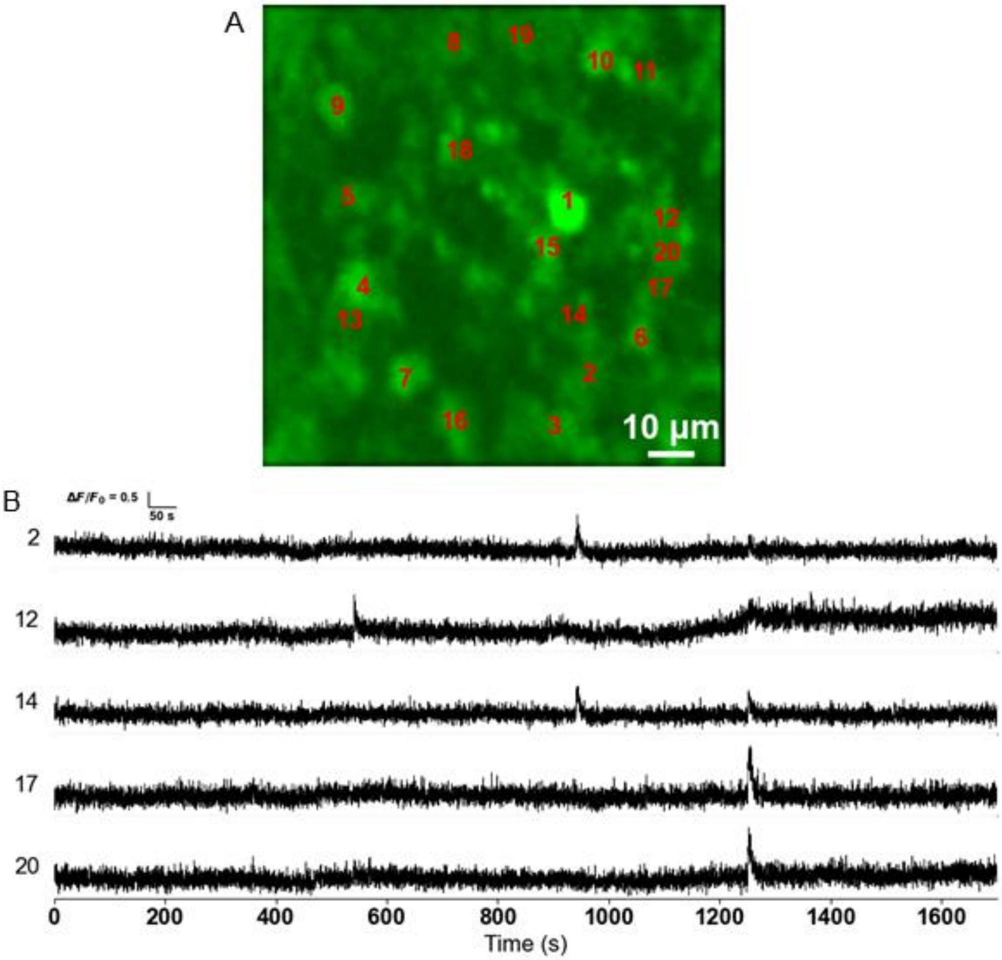


**Supplementary Figure 16**

3P imaging of spontaneous activity in GCaMP8s labelled neurons in DG area of the hippocampus at depth 1725 µm below the pial surface - session #4 from the same FOV as in Fig. 2F and Supplementary Fig. 14-15.

A. Neuronal soma during functional calcium imaging.

B. Fluorescence traces from the active neurons within the DG.

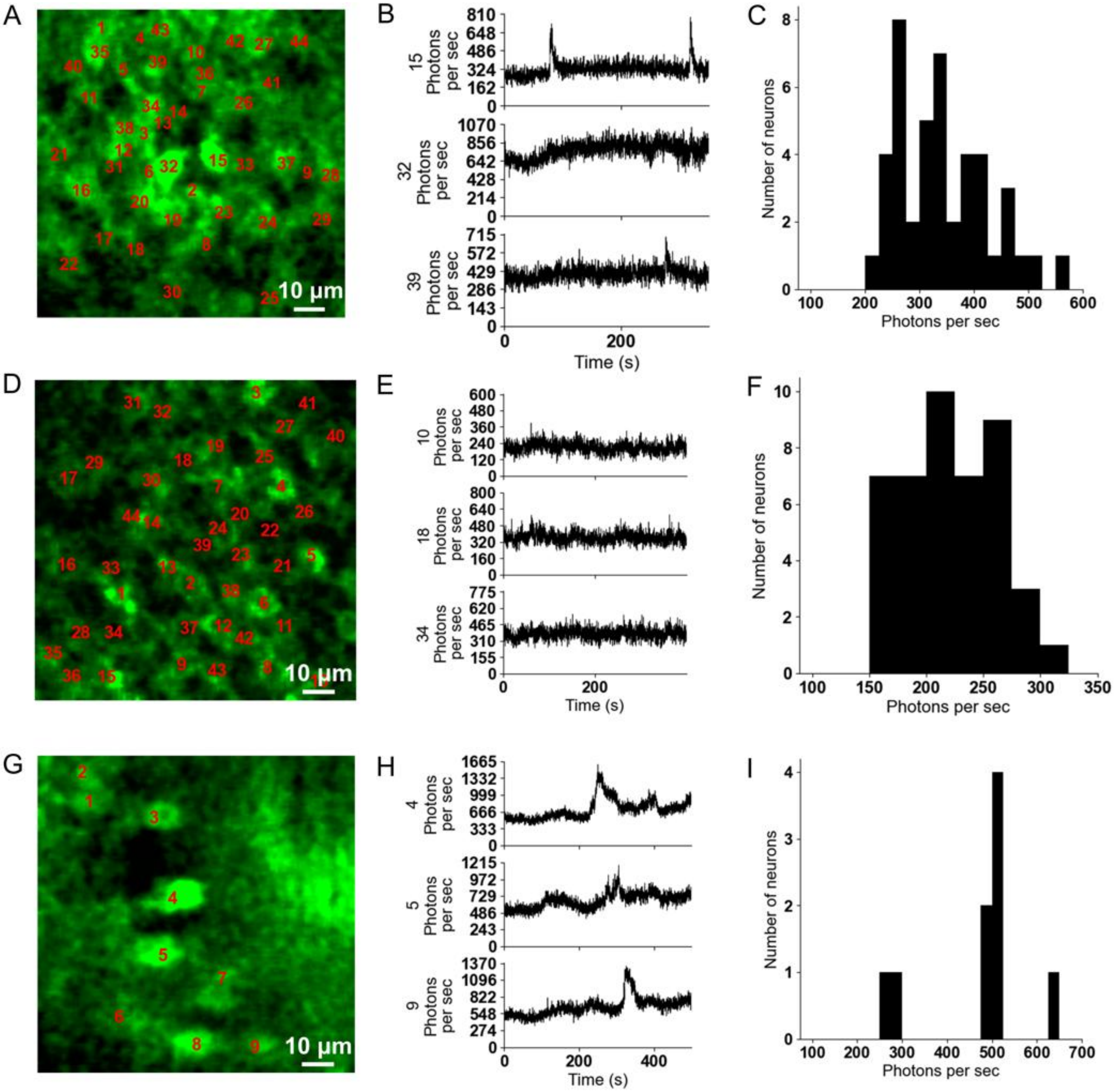


**Supplementary Figure 17**

Photon counts/neuron/s at DG 1725 μm, DG 1820 μm and Cg 2000 μm below the pial surface.

A. Neuronal soma during functional calcium imaging at 1725 μm depth, with DG neurons indicated within the field of view.

B. Representative fluorescence traces from selected neurons within the DG showing spontaneous calcium activity. Y-axis shows the photon counts/neuron/s.

C. A histogram of the baseline photon counts/neuron/s for all neurons indicated in A.

D. E. F. Just like the first row, DG recording at depth 1820 μm, and G. H. I. for Cg recording at depth 2000 μm.

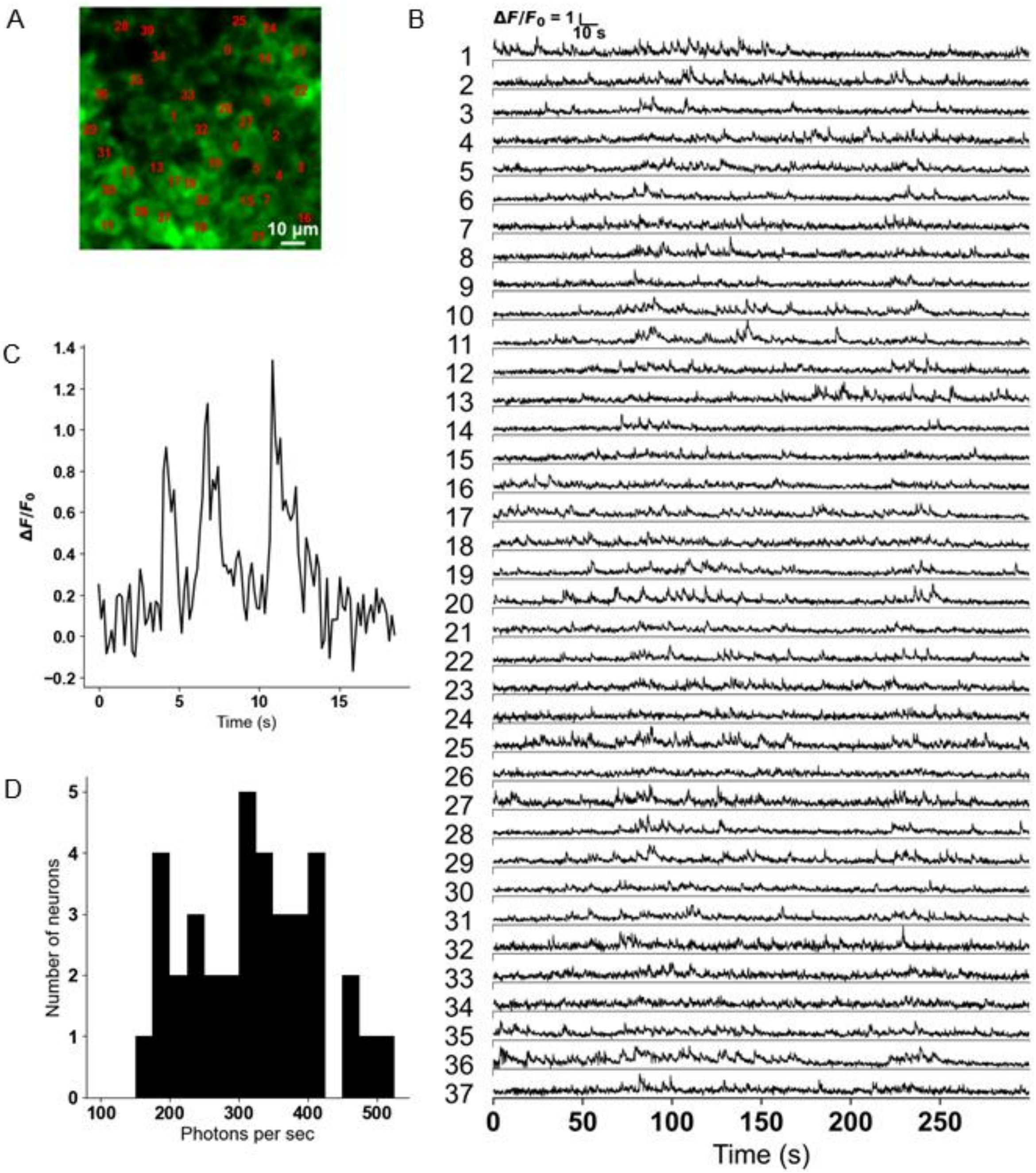


**Supplementary Figure 18**

3P imaging of spontaneous activity in GCaMP8s-labelled neurons in CA1 area of the hippocampus at depth 1090 μm from the brain surface. Repetition rate of laser is 100 kHz, and the frame rate is 6.18 Hz.

A. Neuronal soma during functional calcium imaging.

B. Fluorescence traces from the indicated neurons within the CA1 area.

C. Example of single Ca-transient detection.

D. A histogram of the baseline photon counts/neuron/s for all neurons indicated in A.

**Supplementary Table 1: Imaging parameters (pulse width ~ 60 fs and NA ~ 0.55 for all figures)**

| Figure | Indicator | FOV (µm) | FOV (pixels) | Power Level | | Rep rate | Wavelength (nm) |
|---|---|---|---|---|---|---|---|
| | | | | Depth (µm) | Power (mW) | | |
| Fig 1 Supp Figs. 3-5 | Qdot-605 | 400 x 400 | 400 x 400 | 0-750 | 0.2-2.7 | 100 kHz | 1332 |
| | | | | 750-940 | 2.7-9.1 | | |
| | | | | 940-1400 | 9.1-18.4 | | |
| | | | | 1400-2000 | 18.4-55.8 | | |
| | | | | 2000-2410 | 55.8-94 | | |
| | | | | 2410-2600 | 94 | | |
| Supp Fig. 6 | Fluorescein | 200 x 200 | 400 x 400 | 0-600 | 0.3-2.2 | 100 kHz | 1300 |
| | | | | 600-1300 | 2.2-18.2 | | |
| | | | | 1300-1700 | 18.2-51 | | |
| | | | | 1700-2000 | 51-101 | | |
| | | | | 2000-2200 | 101 | | |
| Fig 2 Supp Fig. 8 | GCaMP8s | 200 x 200 | 400 x 400 | 0-600 | 0.9-7.3 | 500 kHz | 1300 |
| | | | | 600-950 | 7.3-34 | 500 kHz | |
| | | | | 950-1700 | 6.8-68.6 | 100 kHz | |
| | | | | 1700-1800 | 68.6-98 | 100 kHz | |
| | | | | 1800-2000 | 98 | 100 kHz | |
| Supp Fig. 9 | GCaMP8s | 200 x 200 | 400 x 400 | 0-750 | 0.4-7.8 | 500 kHz | 1300 |
| | | | | 750-950 | 7.8-36.8 | 500 kHz | |
| | | | | 950-1500 | 7.4-42.5 | 100 kHz | |
| | | | | 1500-1800 | 42.5-107 | 100 kHz | |
| | | | | 1800-1950 | 107 | 100 kHz | |
| Supp Fig. 10 | GCaMP8s | 400 x 400 | 400 x 400 | 0-1300 | 0.9-101 | 500 kHz | 1300 |
| | | | | 1300-1900 | 20.2-76.5 | 100 kHz | |
| | | | | 1950-2350 | 76.5-101 | 100 kHz | |
| Supp Fig. 11 | GCaMP8s | 400 x 400 | 400 x 400 | 0-900 | 0.9-15.9 | 500 kHz | 1300 |
| | | | | 900-1500 | 15.9-101 | 500 kHz | |
| | | | | 1500-2200 | 25.4-101 | 100 kHz | |
| Supp Fig. 12 | GCaMP8s+ Texas Red | 400 x 400 | 400 x 400 | 0-1500 | 0.47-111 | 500 kHz | 1332 |
| | | | | 1500-1900 | 59.1-99 | 100 kHz | |
| | | | | 1900-2300 | 99 | 100 kHz | |
| Supp Fig. 13 | GCaMP8s | 100 x 100 | 120 x 120 | 1600 | 90.6 | 100 kHz | 1300 |
| Supp Figs. 14-16, 17A-C | GCaMP8s | 100 x 100 | 120 x 120 | 1725 | 107 | 100 kHz | 1300 |
| Supp Fig. 17D-F | GCaMP8s | 100 x 100 | 120 x 120 | 1820 | 107 | 100 kHz | 1300 |
| Supp Fig. 17G-I | GCaMP8s | 100 x 100 | 120 x 120 | 2000 | 107 | 100 kHz | 1300 |
| Supp Fig. 18 | GCaMP8s | 100 x 100 | 120 x 120 | 1090 | 13.8 | 100 kHz | 1300 |

**Supplementary Video List**

1 Qdot_stack
2 Qdot_3D_perspectives
3 Fluorescein_stack
4 DG_stack_mouse_1
5 DG_stack_mouse_2
6 CG_stack_upto_2350
7 CG_stack_upto_2100
8 CG_dual_color_texas_red_and_GCaMP8s
9 DG_activity_depth_1600_raw_registered
10 DG_activity_depth_1600_filtered
11 DG_activity_depth_1725_session_01_raw_registered
12 DG_activity_depth_1725_session_01_filtered
13 DG_activity_depth_1725_session_02_raw_registered
14 DG_activity_depth_1725_session_02_filtered
15 DG_activity_depth_1725_session_03_raw_registered
16 DG_activity_depth_1725_session_03_filtered
17 DG_activity_depth_1725_session_04_raw_registered
18 DG_activity_depth_1725_session_04_filtered
19 CG_activity_depth_2000_raw_registered
20 CG_activity_depth_2000_filtered

Video files are available at https://github.com/monzilur/ultra_deep_3PM
